# Morphological Study of Left–right Head Asymmetry in *Doubledaya bucculenta* (Coleoptera: Erotylidae: Languriinae)

**DOI:** 10.1101/2024.04.05.587748

**Authors:** Hiroki Oda, Taro Nakamura, Wataru Toki, Teruyuki Niimi

## Abstract

Left–right asymmetry in paired organs is well-documented across various species, including the claws of fiddler crabs and snail-eating snakes’ dentition. However, the mechanisms underlying these asymmetries remain largely elusive. This study investigates *Doubledaya bucculenta* (Coleoptera: Erotylidae), a lizard beetle species known for pronounced left-sided asymmetry in adult female mandible and gena. Given that insect mouthparts comprise multiple functionally significant appendages, we aimed to clarify the degree of asymmetry extending beyond the mandibles and genae. Phenotypic morphology was assessed through trait measurement and asymmetry index calculations. Our detailed morphometric analyses revealed left-longer asymmetry not only in mandibles and genae but also in maxillae and labium. Notably, the degree of asymmetry in other mouthparts was generally less pronounced compared to outer mandibles, suggesting a potential influence of left mandible development on other mouthparts. Additionally, male mandibles exhibited region-specific asymmetry, potentially indicative of constrained evolutionary adaptations. This study enhances a comprehensive understanding of adult phenotype morphology and offers insights into the developmental basis of asymmetrical mouthparts.

## Introduction

Investigations into left–right asymmetry, particularly employing model organisms like mice and *Drosophila melanogaster*, have focused on the developmental dynamics of singular organs such as the heart and digestive tracts. Early developmental investigations suggest that ciliary motion-induced fluid flow and myosin ID-dependent cellular processes contribute to these asymmetries (reviewed in Nakamura and Hamada, 2012; Okumura et al., 2008). In contrast, less is known about the molecular underpinnings of left–right asymmetry in paired organs, such as insect mandibles and wings, despite many species exhibiting such traits. Instances of such asymmetry are evident in the claws of fiddler crabs (reviewed in Palmer, 2004), the dentition of snail-eating snakes (Hoso et al., 2007), the wings serving as acoustic organs in crickets (Masaki et al., 1987), and the mandibles of genera *Glischrochilus* (Lee et al., 2020), *Manticora* (Oberprieler and Arndt, 2000) and *Oxyporus* (Hanley, 2001).

*Doubledaya bucculenta* (Coleoptera: Erotylidae: Languriinae), a lizard beetle with a remarkably developed left side of the head, presents a unique model for studying left–right asymmetry in paired organs (Lewis, 1884) (Fig. 1). Female adults use their mandibles to bore holes into the internodes of dead bamboo (*Pleioblastus simonii*, *P. chino*, *P. linearis* and *Semiarundinaria* sp.) and lay an egg in the cavities, with morphological adaptations presumed crucial for this behavior (Toki and Togashi, 2011, 2013; Toki and Hosoya, 2012). Larvae move freely within a single internode to propagate yeast *Wickerhamomyces anomalus* and feed and grow (Toki and Togashi, 2011; Toki et al., 2012, 2013). In addition to *D. bucculenta*, other species exhibiting left–right asymmetry in their mouthparts are known to play important roles in their survival and reproduction. For example, cichlids, which are scale-eating fish (Hori, 1993), larvae of water scavengers preying on right-handed snails (Inoda et al., 2003), snail-eating snakes (Hoso et al., 2007), and larvae of the stag beetle chewing wood (Okada et al., 2008), are believed to enhance feeding efficiency through their left–right asymmetrical mouthparts.

**Figure 1.**
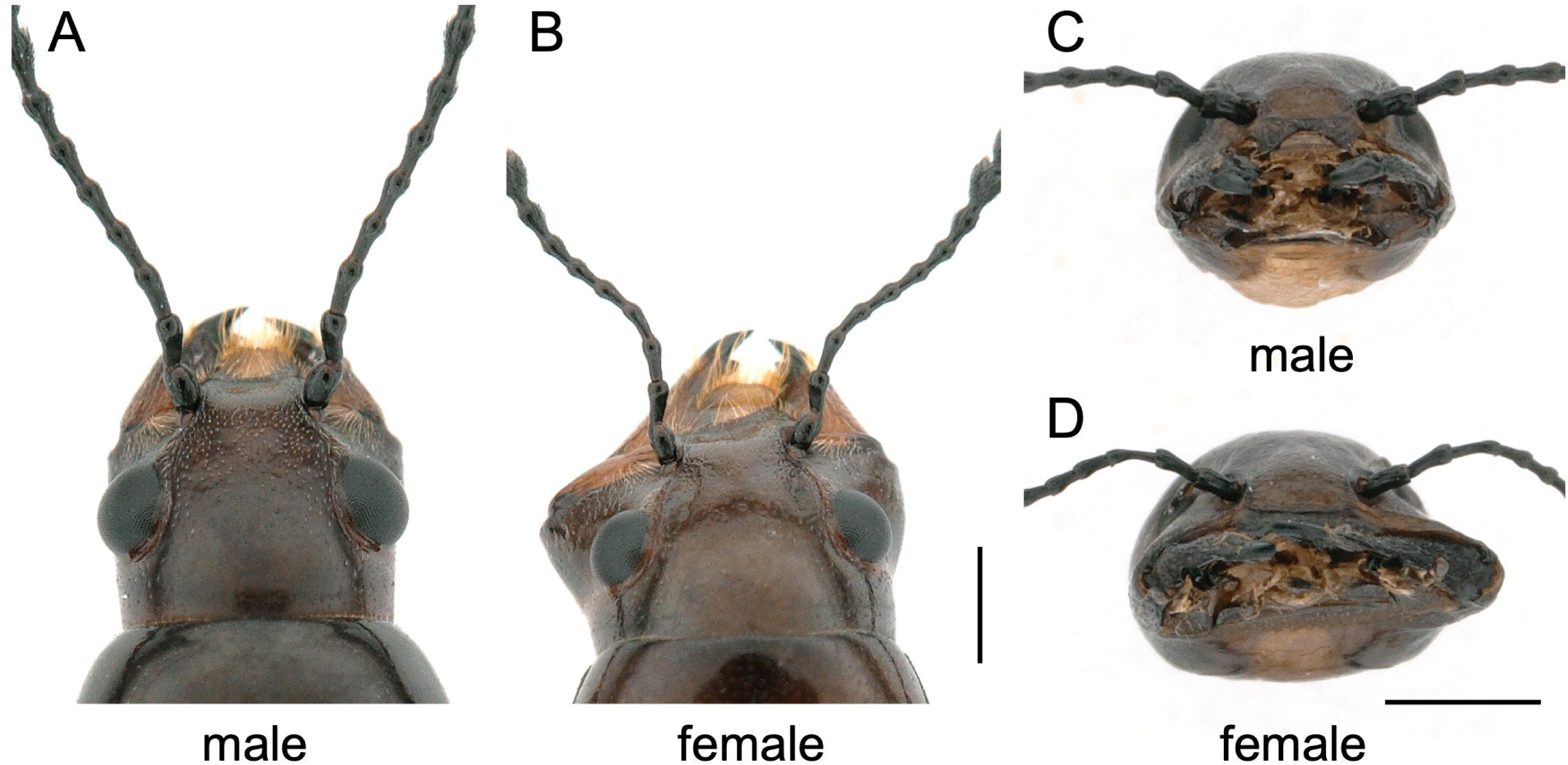
Adult *D. bucculenta* heads. (A) Dorsal view of adult male head; (B) Dorsal view of adult female head; (C) Frontal view of adult male head; (D) Frontal view of adult female head; scales in (A), (B) and (C), (D) are different. The scale bar is 1 mm.

In this study, we focused on the mouthparts of adult *D. bucculenta* and analyzed the left–right asymmetry through morphometric measurements. This insect holds promise as a model for investigating left–right asymmetry in paired organs due to its stable collection from the field and the establishment of an artificial breeding system in previous research (Toki et al., 2012). While previous studies have indicated left–right asymmetry in the gena and mandible (Toki and Togashi, 2011), a comprehensive understanding of detailed morphology and potential asymmetry in other mouthparts remain elusive. Given that the insect mouthparts, including those of *D. bucculenta*, comprise several functional appendages and intricate interplays between form and function, we hypothesize that asymmetries extend beyond the mandibles to other mouthparts in adults. Understanding the phenotypic morphology of mouthparts is crucial for elucidating their functional roles because the morphology of insect mouthparts closely correlates with their feeding behavior and lifestyle (Snodgrass, 1935). To unravel left–right asymmetry in each mouthpart of adult *D. bucculenta*, we assess the degree of asymmetry by calculating the asymmetry index. This study presents morphological traits of the adult phenotype and is expected to contribute to the understanding of the developmental mechanism underlying left–right asymmetry in paired organs.

## Materials and Methods

### Collection of experimental insects

Adults and larvae of *D. bucculenta* were collected from standing dead *P. simonii* in Nishihirose, Toyota, Aichi Prefecture, Japan (35°15’ N, 137°22’ E) between October and December 2021. The bamboo grew naturally in the area, which was neither private property nor a protected area, and no endangered or protected species were distributed there. Adults and larvae were individually collected into 2 ml tubes (Eppendorf) equipped with air holes. Species identification was conducted by comparing morphological characteristics with previously published data (Toki and Togashi, 2011).

### Breeding of D. bucculenta

The breeding method for *D. bucculenta* larvae followed that outlined by Toki et al. (2012). In brief, larvae were placed on 3.9% potato dextrose agar (PDA) medium (Nissui, 05709) without antibiotics, fostering the growth of the symbiotic yeast *W. anomalus*. Larvae were allowed to feed on the yeast spread over to the PDA medium and were transferred to fresh yeast-coated PDA medium every 2–3 weeks. To prevent eclosion failure resulting from adherence to the moist medium, the petri dish was inverted postpupation.

### Dissection of the mouthparts

The head of an adult *D. bucculenta* was dissected by cutting between the head and pronotum with a razor blade. Subsequently, the head was immersed in 0.5 ml of 10% NaOH solution at 60°C for 3 h to soften the mouthparts and the head connective tissues. Following this, the head was rinsed thrice with distilled water using a Pasteur pipette. The mandibles, maxillae, and labium were carefully excised using precision dissecting tweezers (INOX) and air-dried for subsequent morphometric analysis.

### Photography of mouthparts and head

Each dissected mouthpart and head was captured using a digital camera (WRAYMER, FLOYD-2A) mounted on a stereomicroscope (Leica, S9 D) (Fig. 2). Ensuring precise morphometric analysis necessitated adjusting the sample’s position and inclination on the microscope stage to render the three-dimensional morphology as a two-dimensional image. Specific imaging conditions were tailored for each structure to optimize visibility and detail.

**Figure 2.**
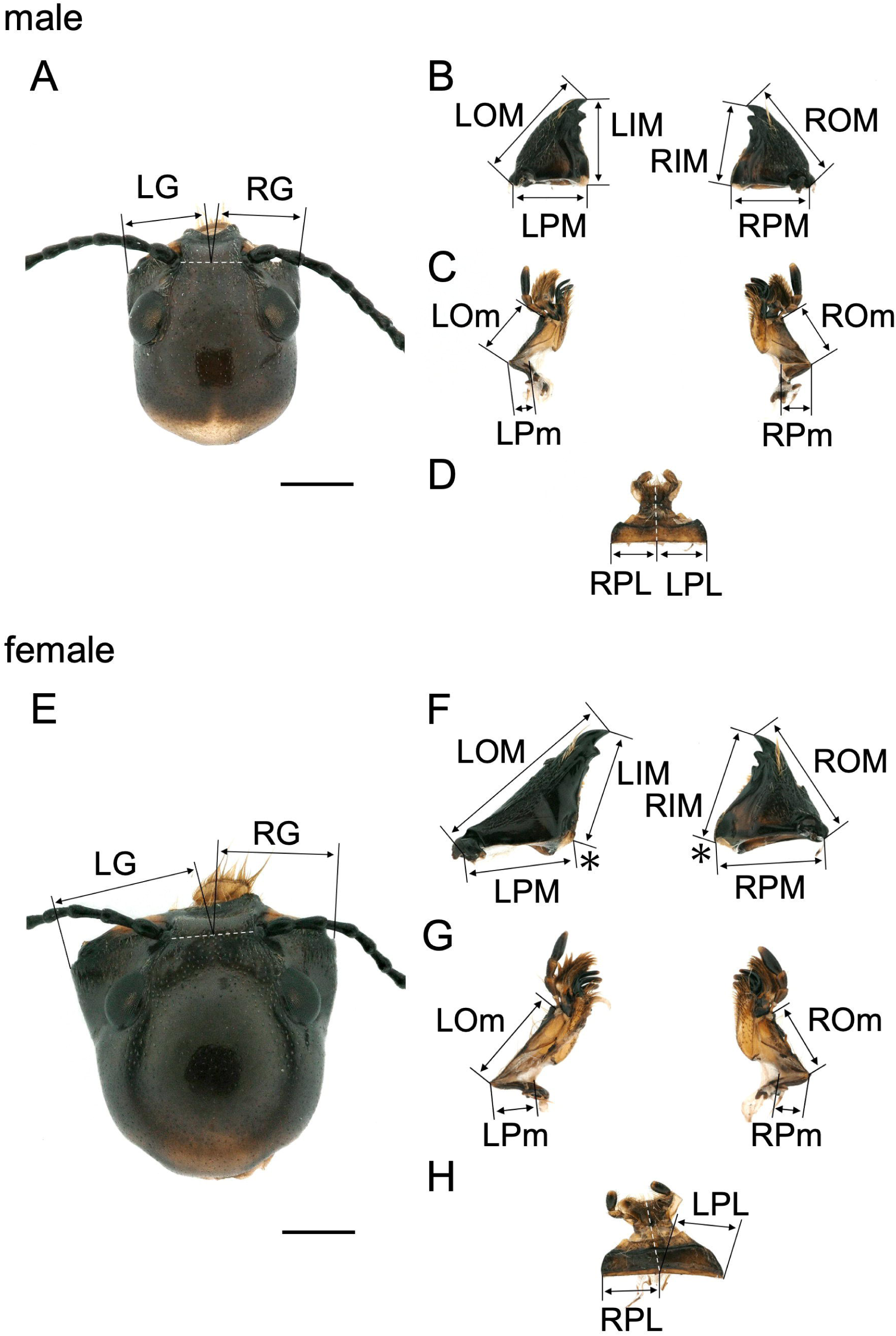
Mouthparts and head traits of adult *D. bucculenta*. (A) Adult male head; (B) Adult male mandibles; (C) Adult male maxillae; (D) Adult male labium; (E) Adult female head; (F) Adult female mandibles; (G) Adult female maxillae; (H) Adult female labium; scale bar is 1 mm. LG, left gena; RG, right gena; LOM, left outer mandible; LIM, left inner mandible; LPM, left proximal mandible; ROM, right outer mandible; RIM, right inner mandible; RPM, right proximal mandible; LOm, left outer maxilla; LIm, left inner maxilla; LPm, left proximal maxilla; ROm, right outer maxilla; RIm, right inner maxilla; RPm, right proximal maxilla; LPL, left proximal labium; RPL, right proximal labium. *The angle between the inner and proximal sides of female mandible.

#### Gena

The head was dorsal side up on a petri dish. Using a kneaded eraser, the angle was adjusted to ensure symmetrical positioning of the compound eyes and centered illumination from the microscope at the base of the antennae. The entire head, with the gena in focus was photographed (Figs. 2 A and 2E).

#### Mandible

After removing any surrounding tissue with a razor blade, the mandible was positioned dorsally on a Kim towel and photographed to capture the entire structure (Figs. 2B and 2F).

#### Maxilla and labium

The maxilla and labium were placed on a slide glass (Matsunami Glass, S1112) and sandwiched between cover glasses (Matsunami Glass, C024601) to facilitate focusing on the entire mouthparts. The maxilla was photographed dorsally (Figs. 2C and 2G), while the labium was captured ventrally (Figs. 2D and 2H).

### Landmark setting and measurement

Landmarks were designated on the gena, mandible, maxilla, and labium of each specimen, and distances between these landmarks were measured using ImageJ software (version 2.9.0). (Fig. 2). Landmarks were selected based on features commonly found on the left and right sides of the mouthparts to ensure accurate measurements despite potential asymmetries. The landmarks and measured areas were as follows.

#### Gena

A line drawn from the midpoint of the line segment connecting the base of the antennae to the projection of the gena. The lengths of the left and right line segments were measured (Figs. 2A and 2E).

#### Mandible

Three points were taken at the tip of the external tooth, the joint associated with the head, and the base of the internal tooth, and the line segments connected were defined as outer, inner, and proximal, respectively, and the length of each line segment was measured (Figs. 2B and 2F).

#### Maxilla

Three points were identified at the base of the maxillary palp and at both ends of its attachment to the head. The length of each line segment, defined as outer and proximal, respectively, was measured (Figs. 2C and 2G).

#### Labium

The central line was extended to the point of attachment to the head, and line segments were drawn from that point to the left and right ends. These segments were considered proximal, and the lengths of the left and right line segments were measured (Figs. 2D and 2H).

Furthermore, the asymmetry index (AI), as defined by Toki and Togashi (2011), was calculated to assess the degree of left–right asymmetry. AI was calculated as {(R − L) × 100} / (R + L), where R and L represented the lengths of the right and left sides of the trait, respectively. AI values ranged from −100 to 100, with 0 indicating symmetrical lengths, negative values indicating a longer left side, and positive values indicating a longer right side.

### Statistical analysis

Welch’s t-test was utilized to compare measurements between left and right (α = 0.05). The distribution of AI values was assumed to be normal, and a bootstrap method (1,000 samples) was employed to calculate a 95% confidence interval (CI) for each AI value. A 95% CI excluding 0 indicated significant left–right bias, while an inclusion suggested symmetry. The analyses were performed using R software (version 4.1.0), with the “simpleboot” (version 1.1-7) and “boot” (version 1.3-28) packages facilitating the bootstrap analysis.

## Results

### Sample collection

We collected a total of 80 specimens (32 male and 35 female adults, and 13 larvae) of *D. bucculenta* from the designated research area between October and December 2021. For morphometric analysis, 30 individuals of each sex were randomly selected from the specimens collected and previously stored in our laboratory.

### Morphological analysis of mouthparts and genae

To elucidate the morphological characteristics of the left and right mouthparts and genae, we measured the distance between predefined landmarks. We focused on the mouthparts involved in perforation and feeding, namely mandibles, maxillae, and labium in this study. Care was taken to position the samples at appropriate angles under the stereomicroscope to minimize potential measurement errors arising from the 3D structure being represented in 2D images. We compared the lengths of each mouthpart and genae on the left and right sides (Fig. 3) and analyzed the frequency distribution of the AI based on the number of individuals (Fig. 4). The mean ± standard deviation, range, and 95% CI for the AI of each trait were summarized in Tables 1 and 2.

**Figure 3.**
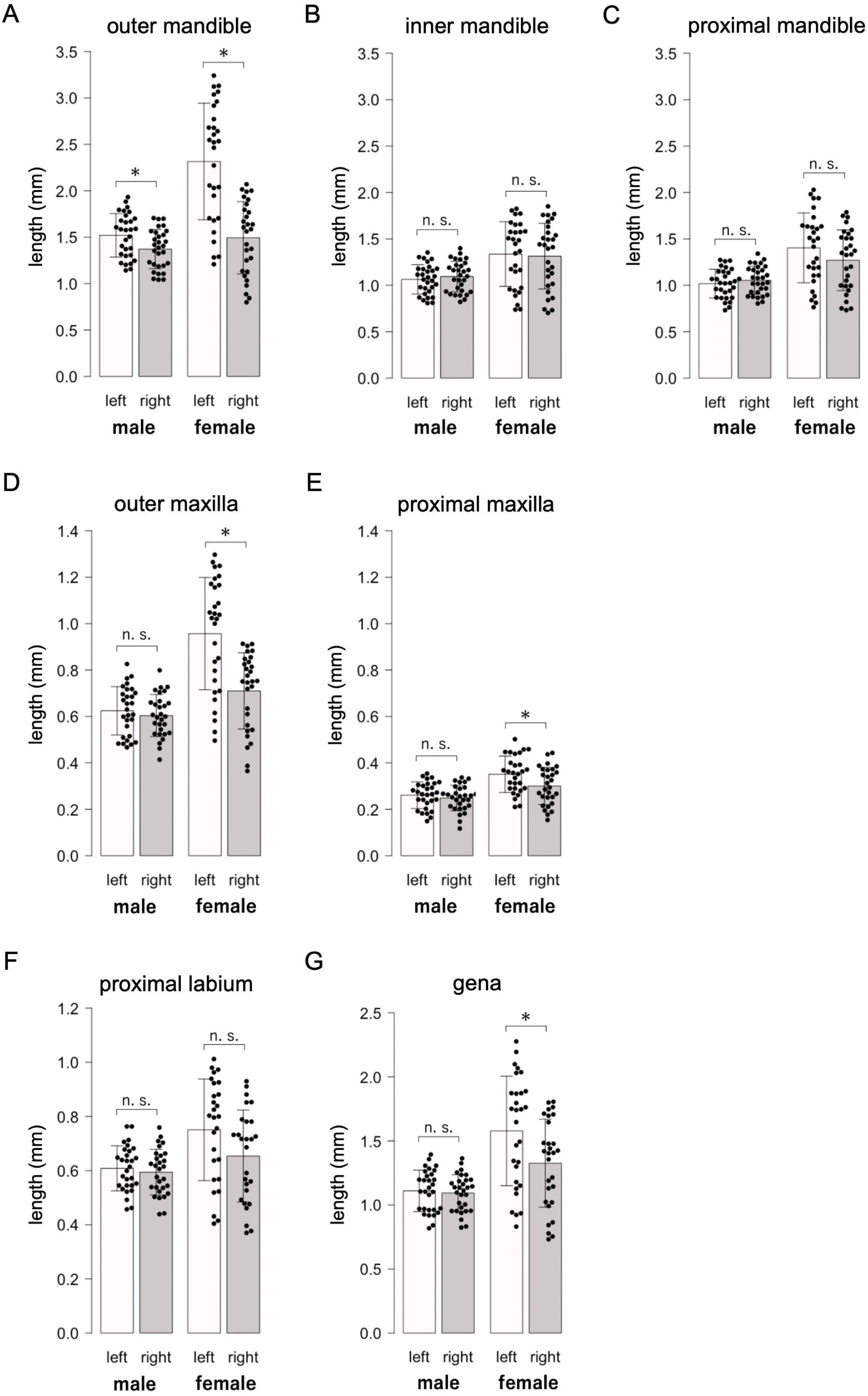
Length of each part of the mouthparts and gena in adult *D. bucculenta*. Left side of each figure is male, right side is female; (A) Outer mandible; (B) Inner mandible; (C) Proximal mandible; (D) Outer maxilla; (E) Proximal maxilla; (F) Proximal labium; (G) Gena; comparisons between the two groups were made by Welch’s t-test. Values are the mean ± SD. *P < 0.05, n. s. = not significant.

**Figure 4.**
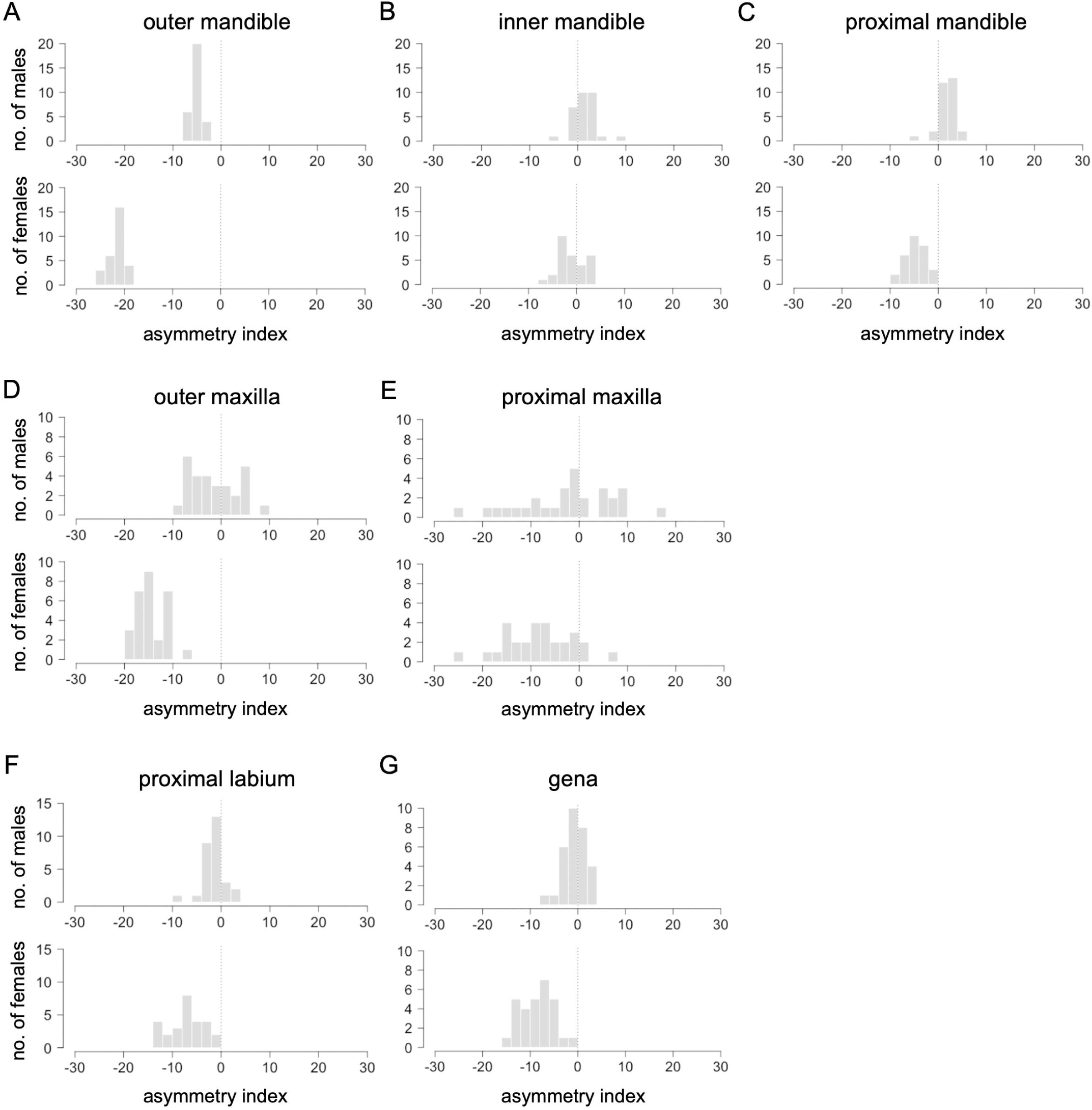
Distribution of asymmetry index (AI) of the mouthparts and gena in adult *D. bucculenta*. Upper side of each figure is male, lower side is female; (A) Outer mandible; (B) Inner mandible; (C) Proximal mandible; (D) Outer maxilla; (E) Proximal maxilla; (F) Proximal labium; (G) Gena. AI is calculated as {(R − L) × 100} / (R + L), where R and L are the lengths of the right and left sides of the trait, respectively.

**Table 1.** Length of each part of the mouthparts and gena in adult *D. bucculenta*. *n* = no. examined.

**Table 2.** Asymmetry index (AI) values of the mouthparts and gena in adult *D. bucculenta*. *n* = no. examined. AI value is calculated as {(R − L) × 100} / (R + L), where R and L are the lengths of the right and left sides of the trait, respectively. AI value = mean ± SE. 95% confidence interval (CI) of mean was obtained by a bootstrap method.

Mandibles analysis revealed a significant difference in the outer length of the left mandible compared to the right in both sexes (males: t_58_ = 2.572, P = 0.013; females: t_56_ = 5.983, P < 0.001) (Fig. 3A). However, no significant difference was identified in the inner or proximal length between the left and right mandibles in either sex (males: inner, t_58_ = −0.727, P = 0.470; proximal, t_58_ = −0.906, P = 0.369; females: inner, t_56_ = 0.247, P = 0.806; proximal, t_56_ = 1.437, P = 0.156) (Figs. 3B and 3C). The AI frequency distribution indicated a left-longer asymmetry in males for the outer mandibles, contrasting with a right-longer asymmetry for the inner and proximal parts. A left-longer asymmetry was observed in females across all mandible sections (Figs. 4A–4C). The 95% CI of AI confirmed these asymmetries except for the inner mandibles in females (Table 2). In this trait, the 95% CI indicated symmetry (Table 2).

The analysis of the maxillae revealed significant differences in the lengths between the left and right outer and proximal maxillae in females (outer, t_56_ = 4.545, P < 0.001; proximal, t_56_ = 2.419, P = 0.019), but not in males (outer, t_56_ = 0.813, P = 0.420; proximal, t_56_ = 0.835, P = 0.407) (Figs. 3D and 3E).

The AI frequency distribution and the 95% CI indicated left-longer asymmetry in females for the outer and proximal maxillae, while males exhibited symmetry in these traits (Figs. 4D and 4E; Table 2).

Labium analysis found no significant difference in the length between the left and right proximal labium in both sexes (males: t_56_ = 0.651, P = 0.518; females: t_52_ = 1.992, P = 0.052) (Fig. 3F). However, the AI frequency distribution showed left-longer asymmetry for the proximal labium in both sexes (Fig. 4F), supported by the 95% CI not including 0 (Table 2).

Genae analysis demonstrated a significant difference in the lengths between the left and right genae in females (t_56_ = 2.474, P = 0.017), but not in males (t_58_ = 0.424, P = 0.673) (Fig. 3G). The AI frequency distribution indicated a left-longer asymmetry in females, with the 95% CI confirming this finding, while males exhibited symmetry in this trait (Fig. 4G, Table 2).

Overall, our results revealed left–right asymmetry in mouthpart traits and genae, with females displaying more traits indicating left-longer asymmetries and exhibiting a greater degree of left-longer asymmetries than males (Fig. 3, Table 2). This was particularly evident in the outer mandibles, where both sexes displayed left-longer asymmetry, with females showing more pronounced differences than other mouthparts traits (Figs. 3 and 4). In female mandibles, the angle between the inner and proximal sides on the left was obtuse, while on the right, it was acute (Fig. 2F). Furthermore, the AI analysis highlighted these asymmetries across various mouthparts and genae, detecting right-longer asymmetries only in males for the inner and proximal mandibles. (Table 2).

## Discussion

This study provides a detailed analysis of the geometric morphology of the mouthparts and head in *D. bucculenta*, revealing notable left–right asymmetries across several components in female adults. The observed asymmetry in mandibles and genae aligns with findings from a previous study (Toki and Togashi, 2011). Additionally, we have identified new left–right asymmetry in other mouthparts, including maxillae and labium. Below, we note the implications of these intriguing asymmetries in each mouthpart.

### Female mandibles asymmetry

We observed a pronounced left-longer asymmetry in the outer mandibles of female *D. bucculenta*, whereas no such asymmetry was observed in the inner or proximal mandibles. This finding suggests that asymmetrical morphogenesis in *D. bucculenta* is site-specific within the mandibles rather than across the entire structure. This notion is supported by the differing angles between the inner and proximal sides of the left and right mandibles. The elongated left outer mandible likely serves a specialized role during bamboo perforation, leveraging its shape for deeper penetration into the internode. This functional adaptation is consistent with previous observations of females using their left mandible for spawning behavior (Toki and Togashi, 2011).

### Male mandibles asymmetry

Left-longer asymmetry in the outer mandibles is common to both sexes. However, males exhibit a lesser asymmetry in the outer mandibles compared to females. This difference may stem from the reduced need for mandible use in males, primarily for emerging from the bamboo. In related bamboo-using species such as *D. ruficollis*, *D. sinuata*, *D. tonkinensis*, and *Oxylanguria acutipennis*, left-longer mandibles are present in both sexes, with males displaying less pronounced asymmetry compared to females (Toki et al., 2016, 2019). This shared phenomenon may represent a characteristic of the ancestral species of *D. bucculenta*.

Interestingly, mandible asymmetry in males has been linked to fighting in male–male competition. Adult males of the genus *Agathidium* possess asymmetrical mandibles, with a horn on the left mandible. Individuals with larger horns have an advantage in the male–male competition for a female (Miller and Wheeler, 2005). In *D. bucculenta*, observations indicate that mandibles are used for fighting between males (Toki and Togashi, 2011), suggesting that mandible asymmetry may play a crucial role in male competition. This developmental constraint on asymmetrical mandibles suggests a potential ancestral trait, warranting further investigation across the Coleoptera family.

### Maxillae and labium asymmetry

The AI analysis also revealed left-longer asymmetry in the outer maxillae and proximal labium, particularly pronounced in females. In Coleoptera, the mouthparts retain ancestral traits, with most possessing biting mandibles and maxillae for feeding (Krenn, 2019). Adult *D. bucculenta* use their maxillae to consume food on bamboo surfaces (Toki and Togashi, 2011). Maxillae and labium are not thought to be involved in bamboo perforation. The mouthparts, comprising head appendages, are serially homologous, and share partially overlapping formation mechanisms (Snodgrass, 1935). RNAi experiments conducted on adult *Tribolium castaneum* revealed many gene functions in the maxilla and labium, suggesting homology of mouthparts at the developmental patterning level (Angelini et al., 2012). These findings suggest that the asymmetries observed in the maxillae and the labium in *D. bucculenta*, particularly among females, may arise as secondary effects stemming from the pronounced development of left-longer mandibles and genae, rather than direct functional adaptations. This interpretation is further supported by the smaller AI of the maxillae and labium compared to the outer mandibles.

### Left–right asymmetry in the genae

The genae were symmetrical in males, while the left gena was longer than the right gena in females. In adult females exhibiting a prominently protruding left gena, the volume within the head between the left and right sides, indicating a left–right difference in the amount of muscle occupying the interior of the head. In insect morphology, mandibular movement primarily relies on the action of adductor muscles, which anchor to the inner walls of the head. The generation of biting force results from the contraction of these muscles’, culminating to the closure of the mandibles (Snodgrass, 1935). In rove beetles (Coleoptera: Staphylinidae), the adductor muscles are known to attach to the genae among other regions of the inner head wall. Species characterized by robust mandibles typically exhibit well-developed adductor musculature, with a conspicuous enlargement of head regions serving as attachment sites for these muscles (Li et al., 2011). Therefore, the observed asymmetry, particularly the enhanced development of the left gena in adult females, may contribute to amplifying the biting force generated in the mandible. Nondestructive 3D reconstructions of the interior of the head using micro-CT will reveal more details by identifying the amount of muscle responsible for mandible movement and the areas where the muscles attach.

### Relationship between body length and left–right asymmetry in *D. bucculenta*

Previous studies have noted a positive correlation between body size and specific morphological traits in *D. bucculenta*, such as head width, genae size, and mandible length. However, the degree of asymmetry in mandibles and genae appears to be independent of body size (Toki and Togashi, 2011). Furthermore, the AI across various mouthparts typically falls within a narrow range of ±10%, with the notable exception of the proximal maxillae. This pattern implies that the extent of asymmetry in these structures remains remarkably consistent among individuals. In the broader context of coleopteran morphology, certain species that exhibit an allometric relationship between body size and exaggerated traits, such as the enlarged mandibles seen in male stag beetles, which become more pronounced with increased body size—a clear example of sexually dimorphic and male-specific adaptation (Romiti et al., 2015; Chen et al., 2020). Contrary to this, the asymmetry in the mouthparts and genae of *D. bucculenta* develops regardless of sex and body size. This suggests that the degree of asymmetry adapted for the task of piercing through bamboo during emergence does not correlate with the beetle’s body size. The consistent extent of asymmetry across individuals implies that this asymmetrical development is a species-specific adaptation closely linked to the unique life cycle of *D. bucculenta*, spent within the confines of bamboo internodes.

In this study, the number of traits showing significant differences in length between the left and right sides was fewer than those indicating left–right asymmetry based on the AI analysis. One reason for this could be that we did not account for the significant individual differences in body length among *D. bucculenta* adults. Our study offers new insights into the left–right asymmetry of the adult phenotype in *D. bucculenta* and contributes to understanding the morphological traits of mouthparts. Future investigations into unmeasured mouthparts morphology will help determine the extent to which asymmetrical mandible morphology is influenced. We anticipate that this study will contribute to elucidating the genetic and developmental mechanism of left–right asymmetry in paired organs.

## Acknowledgments

We thank Y. Chikami for help with the statistical analysis. We extend our gratitude to the Emerging Model Organisms Facility of NIBB Trans-Scale Biology Center for technical assistance. This work was supported by JSPS KAKENHI Grant Numbers 20K21436 to T. Nakamura and the Sumitomo Foundation to T. Nakamura.

## Author Contributions

H. O., T. Nakamura, and T. Niimi conceived the project. H. O., T. Nakamura, and W. T. collected *D. bucculenta*. H. O. performed all experiments associated with morphometric analysis and statistical analysis. All authors contributed to the initial draft of the paper and provided comments on the manuscript.

## References

Angelini DR, Smith FW, Aspiras AC, Kikuchi M, Jockusch EL (2012) Patterning of the Adult Mandibulate Mouthparts in the Red Flour Beetle, *Tribolium castaneum*. Genetics 190: 639–654

Chen ZY, Hsu Y, Lin CP (2020) Allometry and Fighting Behaviour of a Dimorphic Stag Beetle *Cyclommatus mniszechi* (Coleoptera: Lucanidae). Insects 11: 81

Hanley RS (2001) Mandibular allometry and male dimorphism in a group of obligately mycophagous beetles (Insecta: Coleoptera: Staphylinidae: Oxyporinae). Biol J Lin Soc 72: 451–459

Hori M (1993) Frequency-Dependent Natural Selection in the Handedness of Scale-Eating Cichlid Fish. Science 260: 216–219

Hoso M, Asami T, Hori M (2007) Right-handed snakes: convergent evolution of asymmetry for functional specialization. Biol Lett 3: 169–172

Inoda T, Hirata Y, Kamimura S (2003) Asymmetric Mandibles of Water-Scavenger Larvae Improve Feeding Effectiveness on Right-Handed Snails. Am Nat 162: 811–814

Krenn HW (2019) Insect Mouthparts: Form, Function, and Performance. Springer Press

Lee MH, Lee S, Lee S (2020) Review of the subfamily Cryptarchinae Thomson, 1859 (Coleoptera: Nitidulidae) in Korea (Part I: genus *Glischrochilus* Reitter, 1873 and *Pityophagus* Shuckard, 1839). J Asia Pac Biodivers 13: 349–357

Lewis G (1884) Japanese Languriidae, with Notes on their Habits and External Sexual Structure. J Linn Soc Lond Zool 17: 347–361

Li D, Zhang K, Zhu P, Wu Z, Zhou H (2011) 3D configuration of mandibles and controlling muscles in rove beetles based on micro-CT technique. Anal Bioanal Chem 401: 817–825

Masaki S, Kataoka M, Shirato K, Nakagahara M (1987) Evolutionary differentiation of right and left tegmina in crickets. Evol Biol Orthopteroid Insects Ed Baccio MB: 347–357

Miller KB, Wheeler QD (2005) Asymmetrical male mandibular horns and mating behavior in *Agathidium* Panzer (Coleoptera: Leiodidae). J Nat Hist 39: 779–792

Nakamura T, Hamada H (2012) Left-right patterning: conserved and divergent mechanisms. Development 139: 3257–3262

Oberprieler RG, Arndt E (2000) On the biology of *Manticora* Fabricius (Coleoptera: Carabidae: Cicindelinae), with a description of the larva and taxonomic notes. Tijdschr Entomol 143: 71–89

Okada Y, Fujisawa H, Kimura Y, Hasegawa E (2008) MorphDdependent form of asymmetry in mandibles of the stag beetle *Prosopocoilus inclinatus* (Coleoptera: Lucanidae). Ecol Entomol 33: 684–689

Okumura T, Utsuno H, Kuroda J, Gittenberger E, Asami T, Matsuno K (2008) The Development and Evolution of Left-Right Asymmetry in Invertebrates: Lessons from *Drosophila* and Snails. Dev Dyn 237: 3497–3515

Palmer AR (2004) Symmetry Breaking and the Evolution of Development. Science 306: 828–833

Romiti F, Tini M, Redolfi De Zan L, Chiari S, Zauli A, Carpaneto GM (2015) Exaggerated Allometric Structures in Relation to Demographic and Ecological Parameters in *Lucanus cervus* (Coleoptera: Lucanidae). J Morphol 276: 1193–1204

Snodgrass RE (1935) Principles of Insect Morphology. Cornell University Press

Toki W, Hosoya T (2012) New Host Plant and Southernmost Records of Asymmetric Lizard Beetle *Doubledaya bucculenta* Lewis (Coleoptera, Erotylidae, Languriinae). Elytra, New Series 1: 253–254

Toki W, Togashi K (2011) Exaggerated Asymmetric Head Morphology of Female *Doubledaya bucculenta* (Coleoptera: Erotylidae: Languriinae) and Ovipositional Preference for Bamboo Internodes. Zool Sci 28: 348–354

Toki W, Togashi K (2013) Relationship between Oviposition Site Selection and Mandibular Asymmetry in Two Species of Lizard Beetles, *Anadastus Pulchelloides* Nakane and *Doubledaya bucculenta* Lewis (Coleoptera: Erotylidae: Languriinae). Coleopts Bull 67: 360–367

Toki W, Tanahashi M, Togashi K, Fukatsu T (2012) Fungal Farming in a Non-Social Beetle. PLOS ONE 7: e41893

Toki W, Takahashi Y, Togashi K (2013) Fungal Garden Making inside Bamboos by a Non-Social Fungus-Growing Beetle. PLOS ONE 8: e79515

Toki W, Pham HT, Togashi K (2016) Relationship Between Mandibular Asymmetry, Oviposition Hole, and Oviposition Substrate Hardness in Two Bamboo-Using Lizard Beetles *Doubledaya tonkinensis* and *D. sinuata* (Coleoptera: Erotylidae: Languriinae). Ann Entomol Soc Am 109: 850–859

Toki W, Matsuo S, Pham HT, Meleng P, Lee C (2019) Heads or tails: exaggerated morphologies in relation to the use of large bamboo internodes in two lizard beetles, *Doubledaya ruficollis* and *Oxylanguria acutipennis* (Coleoptera: Erotylidae: Languriinae). Sci Nat 106: 50

